# Sequence fingerprints distinguish erroneous from correct predictions of Intrinsically Disordered Protein Regions

**DOI:** 10.1101/203547

**Authors:** Konda Mani Saravanan, A Keith Dunker, Sankaran Krishnaswamy

## Abstract

More than sixty prediction methods for intrinsically disordered proteins (IDPs) have been developed over the years, many of which are accessible on the world-wide web. Nearly, all of these predictors give balanced accuracies in the ~65% to ~80% range. Since predictors are not perfect, further studies are required to uncover the role of amino acid residues in native IDP as compared to predicted IDP regions. In the present work, we make use of sequences of 100% predicted IDP regions, false positive disorder predictions, and experimentally determined IDP regions to distinguish the characteristics of native versus predicted IDP regions. A higher occurrence of asparagine is observed in sequences of native IDP regions but not in sequences of false positive predictions of IDP regions. The occurrences of certain combinations of amino acids at the pentapeptide level provide a distinguishing feature in the IDPs with respect to globular proteins. The distinguishing features presented in this paper provide insights into the sequence fingerprints of amino acid residues in experimentally-determined as compared to predicted IDP regions. These observations and additional work along these lines should enable the development of improvements in the accuracy of disorder prediction algorithm.

## INTRODUCTION

Intrinsically disordered proteins (IDPs) and IDP regions lack a fixed tertiary structure, yet are involved in a wide variety of biological functions (Wright & Dyson, 1999; Dunker et al., 2001; Uversky, 2002; Tompa, 2002; Gunasekaran et al., 2003; Dyson & Wright, 2005; Xue, Dunker & Uversky, 2012; Forman-Kay & Mittag, 2013; Oldfield & Dunker, 2014). For example, IDPs and IDP regions play varied functional roles by linking other structural elements and by binding to their own active or allosteric sites to auto-inhibit or activate their own functions Yeon et al., 2016) or by binding to other molecules, such as structured proteins, DNA, RNA, membranes and various small molecules or even ions, thereby forming dynamic complexes that are used for function (Wright & Dyson, 1999; Dunker et al., 2001; Uversky, 2002; Tompa, 2002; Gunasekaran et al., 2003; Dyson & Wright, 2005; Xue, Dunker & Uversky, 2012; Forman-Kay & Mittag, 2013; Oldfield & Dunker, 2014; Yeon et al., 2016). IDPs and IDP regions also play crucial roles in the assembly and functions of complex molecular machines such as the ribosome (peng et al., 2014), the splicesosome (Korneta & Bujnicki, 2012; Coelho Ribeiro Mde et al., 2013), the mediator complex (Tóth-Petróczy et al., 2008) and the β-catenin destruction complex (Xue et al., 2013). More recently IDPs and IDP regions have been shown to be present and likely important for the formation and function of various membrane-less organelles (Uversky et al., 2015; Shmidt & Görlich, 2016).

Given the various and important functions of IDPs and IDP regions, several databases have been developed to aid in the classification and to provide information regarding the distribution of IDPs and IDP regions in the protein universe (Sickmeier et al., 2007; Di Domenico et al., 2012; Oates et al., 2013; Fukuchi et al., 2014; Varadi et al., 2014; Piovesan et al., 2017). Using the information found in these databases and elsewhere, many computational methods have been developed to predict intrinsic disorder from protein sequences (Romero et al., 1997; Dosztányi et al., 2005; Ishida & Kinoshita, 2007; He et al., 2009; Xue et al., 2010; Walsh et al., 2012; Monastyrskyy et al., 2014). Several protein science research groups have analyzed protein intrinsic disorder from the perspective of genomics and proteomics (Dunker et al., 2000; Linding et al., 2003; Ward et al., 2004; Radivojac et al., 2007). Since IDP regions occupy a considerable percentage of residue positions in most viral (Minde, Dunker & Lilley, 2017) and all non-viral proteomes so far investigated (Xue, Dunker & Uversky, 2012; Oates et al., 2013), it has become important to analyze the sequences of these proteins with a variety of computational approaches to uncover the conserved structural and functional features of these proteins (Chen et al., 2006; Saravanan & Selvaraj, 2017).

Based on amino acid composition of IDP regions, an amino acid index (scale) has been constructed helps us to understand about the specific residues that promote order or disorder (Campen et al., 2008). Evolutionary based approaches have also been developed to study the proteins with IDP regions (Brown et al., 2002; Fukuchi et al., 2006; Szalkowski & Anisimova, 2011; Bellay et al., 2011; Brown et al., 2011; Schlessinger et al., 2011; Bannerjee, Chakraborty & De, 2017). It was also shown that the tandem amino acid repeats accumulate in IDP regions and often play vital roles in the evolution of these regions (Dosztányi et al., 2006; Simon & Hancock, 2009).

Pattern and motif analysis of IDP regions in human sequences as well as other sequence analysis approaches applied to all types of organisms has revealed that the functions of IDPs are very common and are length and position dependent (Lobley et al., 2007; Yan et al., 2016; Meszaros, Dosztanyi & Simon, 2012). While amino acid composition has been frequently used to predict protein disordered regions, the alternative of using a reduced amino acid alphabet model has some advantages (Weathers et al., 2004; Shimizu, Hirose, & Noguchi, 2007). Recently, the distribution and cluster analysis of predicted intrinsically disordered protein Pfam domains has been reported, for which 12456 unique Pfam domains have been considered (Williams et al., 2013). Further, these 12456 Pfam domains have been subjected to disorder prediction by using three disorder predictors like VL3 (Obradovic et al., 2003), VSL2B (Obradovic et al., 2005), and VLXT (Romero et al., 2001) and the consensus have been reported. Since Pfam domains are typically assumed to be structured, the finding that some are predicted to be IDP regions has important implications for the analysis of protein structure and function.

The Critical Assessment of Structure Prediction (CASP) experiment included a section for disorder predictors from CASP 5 (2002) to CASP 10 (2012) (Monastyrskyy et al., 2014). While the CASP experiment led to the development of more than 40 online disorder predictors, there has been very little and perhaps no systematic improvement in disorder prediction over this 10-year span (Dunker & Oldfiels, 2015), suggesting that new ideas are needed in this field. Indeed, the error rate for IDP prediction has remained quite high, in the range of 20% or more (Walsh et al., 2012; Peng & Kurgan, 2012; Potenza et al., 2015). Also, the number of predicted IDP residues available for study (Oates et al., 2013), greatly exceeds the number of experimentally determined IDP residues (Sickmeier et al., 2007). Thus, for IDP regions likely involved in especially important biological function and for future efforts to improve the prediction accuracy of IDPs, it would be useful to have additional analytical tools that can help to identify IDP prediction errors. In the present work, a careful computational sequence fingerprinting method has been developed for comparing 100% predicted IDP regions (Williams et al., 2013) with experimentally determined IDP regions (Sickmeier et al., 2007) and with false positive predictions of IDP regions. These comparisons reveal a number of important distinctions between predicted and experimentally determined IDP regions. These distinctions can be used in the short term to identify likely IDP prediction errors and in the long term to develop improved IDP predictors.

## MATERIALS & METHODS

### DATASET

In order to perform sequence analysis on 100% predicted IDP regions, we have used the dataset of 618 unique Pfam mammalian domains containing 100% predicted IDP regions presented in Table 10 of Williams et al., (2013; Tompa et al., 2009). These authors obtained representative pfam seed set protein sequences of human, mouse and chimp proteomes, and subjected to disorder prediction by VL3 (Obradovic et al., 2003), VSL2b (Obradovic et al., 2005), and VLXT (Romero et al., 2001) to construct 100% predicted IDP regions. By using information from the table (Williams et al., 2013), we have extracted predicted IDP regions and compiled a non-redundant set by using the PISCES server (Wang & Dunbrack, 2003). Since, the sequences in other datasets compared with 100% predicted IDP regions share 25% sequence identity, we have used 25% sequence identity to cull the 100% predicted IDP regions. The 25% sequence identity to cull the dataset reduces the number of sequences; we have used 50% sequence identity to construct a total of 378 predicted IDP regions. To compare the predicted IDP regions with native IDP regions, the latest release of the non-redundant dataset from DISPROT6.02 from DISPROT database is used (Sickmeier et al., 2007; Piovesan et al., 2017). We have used both IDPs (fully disordered proteins) and IDRs (Disordered regions) in the present analysis. Further, the disordered proteins in the dataset were grouped into two sets like intracellular and other secreted proteins. This grouping has been done by searching Uniprot database by looking sub-cellular location entry.

Further, to compare the sequence composition of predicted and native intrinsic disordered amino acid residues with the false positive predicted amino acid residues, we have considered a dataset of fully ordered proteins (Xue et al., 2009). There are 554 chains that were derived from the PDB database to include sequences of non-homologous single chain (typically less than 25% sequence identity in whole protein sequence alignments), non-membrane proteins, which had no ligands, no disulfide bonds, and no missing residues, and which were characterized by unit cells with primitive space groups (Toretsky & Wright, 2014; Wu & Fuxreiter, 2016). All the protein chains in the fully ordered dataset are subjected to consensus disorder prediction (Xue et al., 2010). The results are stored in an appropriate format to perform further computations.

In order to compare the sequence separation of 100% predicted IDP regions with ordered regions, we have downloaded the non-redundant sequences of four major structural classes such as all alpha (280 proteins), all beta (174 proteins), alpha+beta (376 proteins) and alpha/beta (147 proteins) respectively from Structural Classification of Proteins (SCOP) database (Murzin et al., 1995).

### COMPOSITION PROFILES OF PREDICTED AND NATIVE IDP SEQUENCES

The difference between the composition of amino acid residues in predicted, native and false positive predicted IDP regions are computed by using the web server “Composition Profiler” at http://www.cprofiler.org (Vacic et al., 2007). The composition profiles of predicted and native IDP regions are presented based on six important amino acid scales like hydrophobicity, bulkiness, flexibility, alpha-helix forming tendency, beta forming tendency and coil forming tendency. The differences of terminal residues (eight residue fragments of carboxy and amino termini) between predicted and native IDP sequences are computed using the above web server. The frequency of occurrences of dipeptides (400 possible dipeptides) in non-redundant 100% predicted IDPs is computed by using in-house PERL program. It should be noted that the dipeptide repeats are reported to have implications in assembly, dynamics and function of membrane less organelles (Lee et al., 2016).

### SEQUENCE SEPARATION OF AMINO ACID RESIDUES IN 100% PREDICTED IDP SEQUENCES

Sequence separation is defined as the distance of separation of two amino acid residues along a sequence. The number of occurrences of all possible amino acid residue pairs (400 possible amino acid pairs) at various sequence separations (from 1 to 12)^68^ is computed for the 100% predicted IDP sequences by using an in-house program. In order to perform a comparative analysis, we have computed the occurrences of amino acid pairs at 12 sequence separations for the four major structural classes of proteins such as all alpha, all beta, alpha+beta and alpha/beta respectively (Harihar & Selvaraj, 2009). The number of occurrences of an amino acid pair at various sequence separations is stored as 20 × 20 matrices. The sequence separation values of 100% predicted IDP regions are correlated with the sequence separation values of proteins in four major structural classes.

### FINGERPRINTING SEQUENCES OF 100% PREDICTED INTRINSIC DISORDER

In order to perform pattern search, we have considered the amino acid scale constructed by Campen et al., (2008). In this scale, they provided the amino acids from order-promoting residues to disorder-promoting residues (WFYIMLVNCTAGRDHQKSEP). Since it has been shown that a minimum of five residues can form an entire alpha helix or a beta strand, we have used a five residue window to search patterns. The top 3 disorder-promoting amino acids are considered such as P, E, S respectively. We have generated pentapeptides (5 residue fragments) by permutation and combination so that they have these three amino acids, for example, PPPES, PPEES, and so on. As the 3 amino acids could be selected in any order, 243 pentapeptides - called as penta3d(P,E,S) - were generated. Then, we increased the number of disorder promoting amino acid residues to four (P,E,S,K) and generated 1024 pentapeptides designated as penta4d(P,E,S,K). For five disorder promoting amino acid residues (PESKQ), there are 3125 pentapeptides denoted as penta5d(P,E,S,K,Q). Totally, the pentaXd (where X=3,4 or 5) combination consists of the total of the sets penta3d, penta4d and penta5d to give 4392 possible pentapeptides. Similarly, we have generated the pentaXo (here X can be 3,4 or 5) combination of 4392 pentapeptides by using the 5 order promoting residues (WFYIM) that is the sum of the sets penta3o(W,F,Y), penta4o(W,F,Y,I) and penta5o(W,F,Y,I,M). We have used string matching programs to find the number of occurrences of the pentaXd and pentaXo sets in the sequences of Disprot database. By keeping the generated peptides as query, we have made pattern search in sequences of 100% predicted IDP, Disprot and globular proteins. The frequency of the occurrences was taken as the actual number of hits obtained in each set with respect to the total possible number of pentapeptides mentioned above in each case. The expected random frequency of the residue combinations was calculated using the product of the residue frequencies found in each of the datasets. The residue frequencies were calculated based on the total occurrences of the chosen set of amino acids in each dataset (for predicted IDP set: 413315, for Disprot set: 163421, for Globular proteins set: 591114). The ratio of the frequency of occurrence with the expected random frequency was used to compute the log-odds for each of the pentapeptide sets. In order to look for features that can further discriminate the disordered regions and since the generated sequence fragments do not contain repeated amino acid residues like PPP, PPPP, EEE, EEEE, SSS, and SSSS respectively, we have made such pattern search and computed log-odds of occurrences in a similar manner as earlier. We have also computed the number of occurrences of palindromes of different length in 100% predicted IDP sequences. Further, we have searched for short hydrophobic stretches of various lengths in the non-redundant set of predicted and native IDP regions by using PERL programs. We considered the residues V, I, L, M, F, W and C as hydrophobic residues during the search of short stretches of hydrophobic residues.

### SECONDARY STRUCTURE PREDICTION IN 100% PREDICTED IDP SEQUENCES

In order to find, how the secondary structure prediction methods perform over the disordered predictors, we have performed consensus secondary structure prediction on the 100% predicted IDP sequences. We have used consensus secondary structure program developed by Sekar group (Gupta et al., 2009). The method thus used for secondary structure predictions make use of 6 different secondary prediction methods to achieve the consensus (King & Sternberg, 1996; Garnier, Gibrat & Robson, 1996; Frishman & Argos, 1997; Guermeur et al., 1999; McGuffin, Bryson & Jones, 2000; Rost & Eyrich, 2001).

## RESULTS

### AMINO ACID COMPOSITION PROFILES OF PREDICTED, NATIVE AND FALSE POSITIVE PREDICTED INTRINSICALLY DISORDERED PROTEIN REGIONS

The amino acid composition difference between predicted and native IDP regions is shown in Figure 1. The amino acid composition of proteins with predicted intrinsic disorder mostly agrees with the findings of Campen et al., (2008). The major differences noted in the amino acid composition of predicted and native intrinsic disorder are presented in **Supplementary Material 1**. The C residue is highly enriched in the sequences of native IDP regions as compared to the predicted. It was already shown that the addition or removal of cysteine residues might affect the structure and dynamics of intrinsically disordered proteins (Kosol et al., 2013) and hence it is necessary to further analyze the occurrence of cysteine residues in different groups of IDPs. The percentage of occurrences of cysteine residues in intracellular proteins, other secreted proteins and ordered proteins is 1.47, 1.59 and 1.21 respectively. Although, the covalent linkage of cysteine residues by forming disulfide bonds are believed as an essential structural feature of numerous proteins, our results reveal that the abundance of cysteine residues in IDPs (intracellular and other secreted proteins) might play a role in disorder but not in disulfide bonds (Fraga et al., 2014).

**Figure 1.**
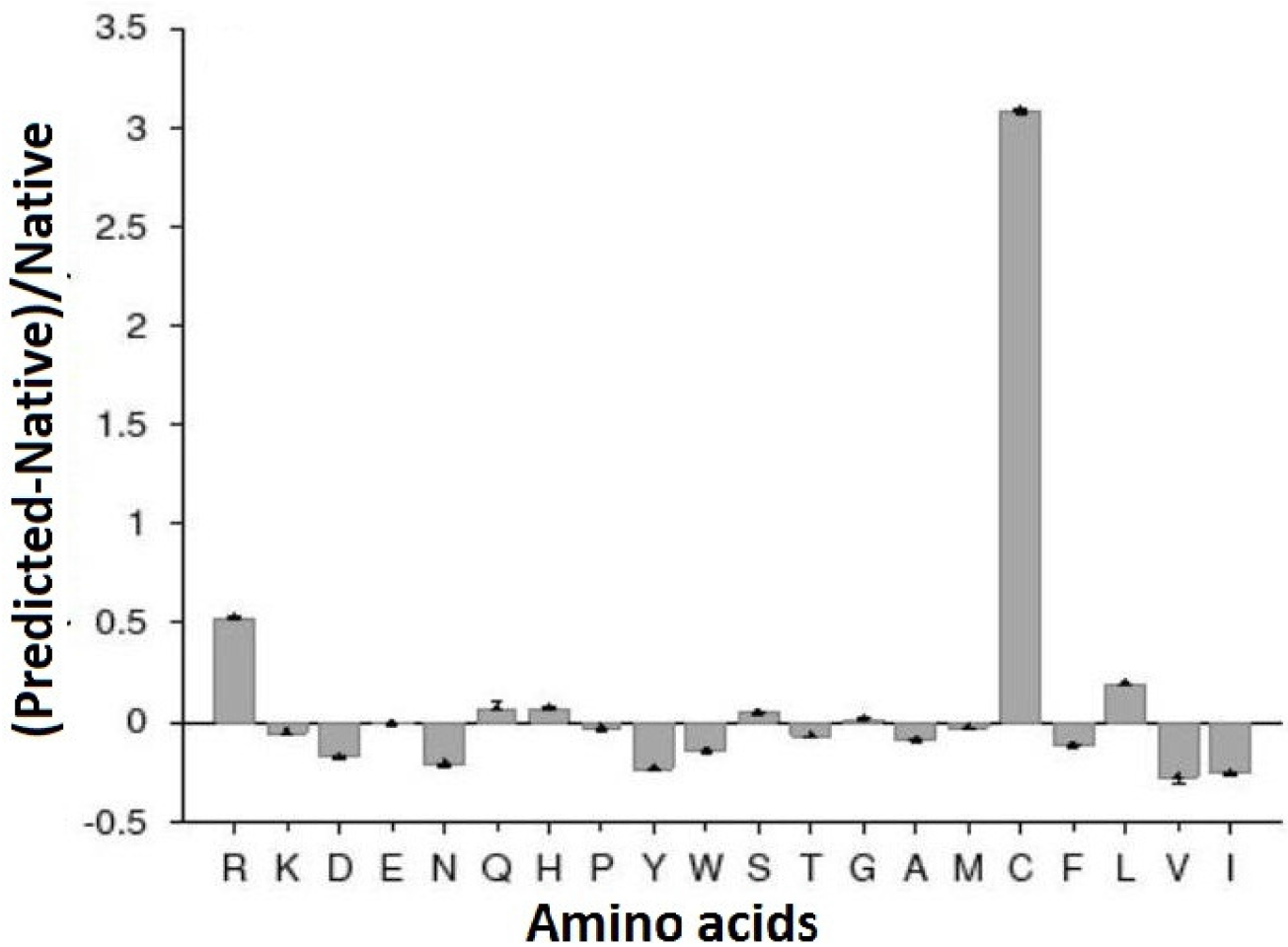
Composition difference profiles of predicted and native IDPs. The positive values indicate that the amino acids are enriched and the negative values indicate that the amino acids are depleted between sequences of predicted and native IDPs. The residue C and positively charged amino acid residue ‘R’ and helix favouring residue ‘H’ and Beta favouring residue ‘L’ is enriched in native IDPs. However, hydrophobic residues (I, V and F), bulky residues (W and Y), flexible residue (D), helix favouring residue ‘A’ and coil favouring residue (N) are depleted in sequences of native IDPs.

The positively charged amino acid residue ‘R’ is enriched in sequences of native IDPs. The hydrophobic residues (I, V and F), bulky residues (W and Y), flexible residue (D) and coil favouring residue (N) are depleted in sequences of native IDPs. The helix favouring residue ‘H’ is enriched in sequences of native IDPs whereas helix favouring residue ‘A’ is depleted. Beta favouring residue ‘L’ is enriched in native IDP sequences. There are no significant composition differences in the residues like E, P, G and M between sequences of predicted and native IDPs.

While comparing the eight residue fragments of amino and carboxy termini of predicted and native IDPs, it is found that the residues like D, G, M, T, W and Y have the same trend in composition differences which is shown in Table 1.

**Table 1.** Amino acid composition differences in amino and carboxy terminii (eight residue fragment) between predicted and native intrinsic disorder.

| S.No | Residue | Differences in amino termini of predicted and native disorder | Differences in carboxy Termini of predicted and native disorder |
| --- | --- | --- | --- |
| 1 | A | Depleted | Depleted |
| 2 | C | Enriched | Enriched |
| 3 | D | Not Significant | Enriched |
| 4 | E | Enriched | Enriched |
| 5 | F | Depleted | Depleted |
| 6 | G | Enriched | Not Significant |
| 7 | H | Not Significant | Not Significant |
| 8 | I | Depleted | Depleted |
| 9 | K | Not Significant | Not Significant |
| 10 | L | Depleted | Depleted |
| 11 | M | Depleted | Not Significant |
| 12 | N | Depleted | Depleted |
| 13 | P | Enriched | Enriched |
| 14 | Q | Enriched | Enriched |
| 15 | R | Enriched | Enriched |
| 16 | S | Enriched | Enriched |
| 17 | T | Depleted | Not Significant |
| 18 | V | Depleted | Depleted |
| 19 | W | Not Significant | Depleted |
| 20 | Y | Not Significant | Depleted |

The residues like C, E, P, Q, R and S are enriched whereas the residues like A, F, I, L, N and V are depleted in the sequences of native IDPs. Only 2.57% of residues are predicted as false positive by the consensus disorder prediction server. While comparing false positive predicted amino acid composition with 100% predicted and native IDP regions amino acid composition, the trend is almost similar as shown in Figure 2. The amino acid residues like N, D, G, S, T are enriched whereas amino acid residues like C, E, H, L, P, Q are depleted. However, small differences are observed between the composition profiles of false positive predicted residues against native and predicted intrinsic disordered residues. The residues like A, I, R are enriched in predicted against false positive IDP sequences whereas there is no significant composition difference in native against false positive predicted IDP sequences.

**Figure 2:**
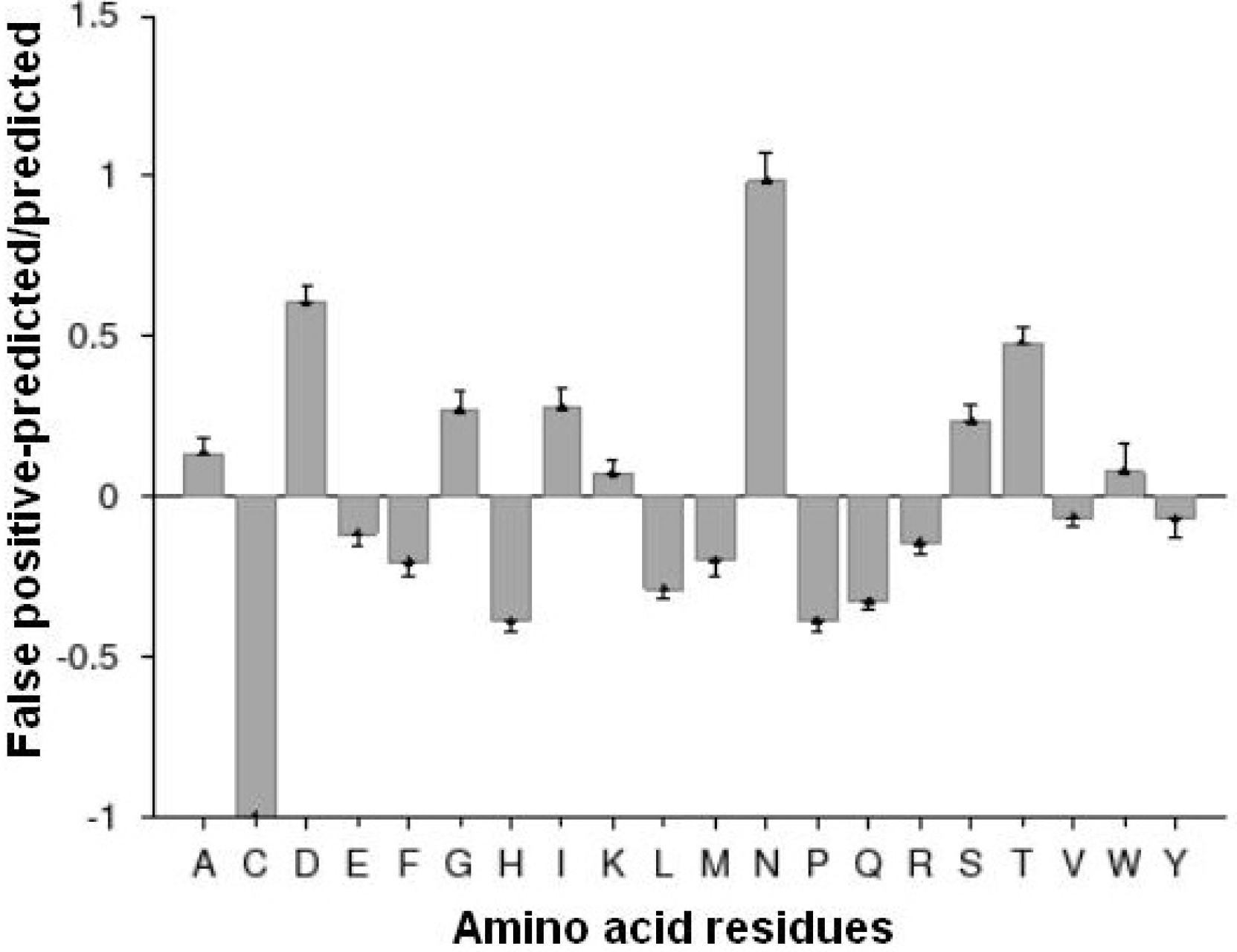
Composition difference profiles of predicted IDPs against false positive predicted IDPs (left) and native IDPs against false positive predicted IDPs (right). The positive values indicate that the amino acids are enriched and the negative values indicate that the amino acids are depleted between sequences of predicted and native IDPs. The residues A, I, R, N, D, G, S, T are enriched in predicted against false positive IDP sequences The amino acid residues C, E, H, L, P, Q, F, V are depleted.

Similarly, the residues like F and V are depleted in predicted against false positive IDP sequences whereas there is no significant composition difference in native against false positive predicted IDP sequences. Further, most of the false positive predictions are of 1-5 amino acids in length. There are a considerable number of false positive predictions of length 6 to 10 and 11 to 15 which is shown in Figure 3.

**Figure 3.**
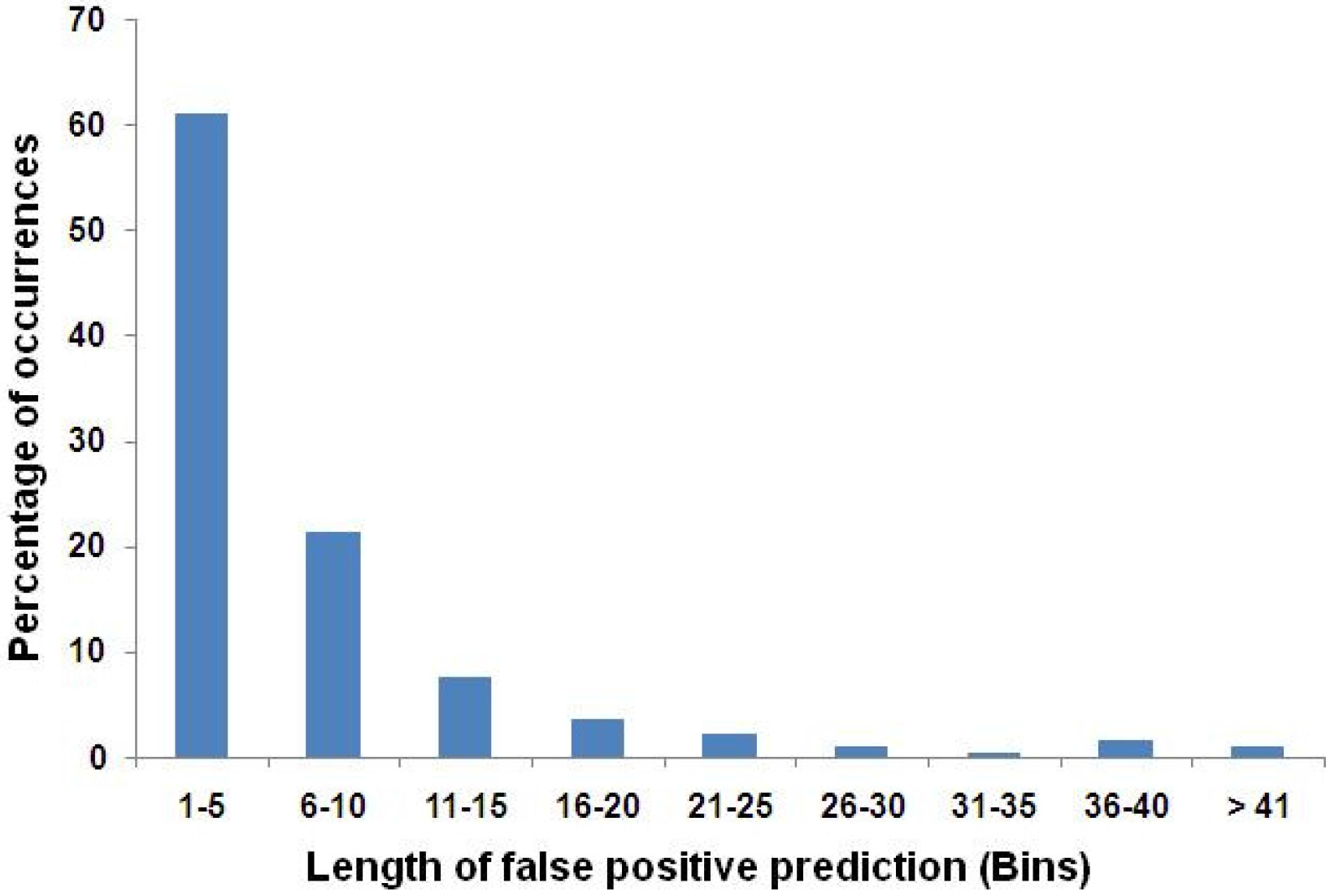
Percentage of occurrences of false positive predicted IDPs of various lengths. The length of the amino acid residues of false positive predicted IDPs are grouped into bins of size 5 to 40. Most of the false positive predictions are of 1-5 amino acids in length. There is a considerable number of false positive predictions of length 6 to 10 and 11 to 15.

### DIPEPTIDE ANALYSIS OF 100% PREDICTED IDP SEQUENCES

The frequency of occurrences of all 400 possible dipeptides in the 100% predicted IDPs is shown as 20×20 grid diagram in Figure 4. The dipeptide EE occurs maximum in number which is shown in red colour in Figure 4. The dipeptides like SS, PP, PG, RR, AA, LE, AE and GP occur in higher frequency. From the figure, it is clear that the first five order promoting (WFYIM) residues have a minimum number of occurrences which are shown as blue grids. The dipeptides containing L as one of the amino acids have a considerable number of occurrences. There are other dipeptides with disorder-promoting residues which occur in considerable numbers as shown in white colour. The maximum occurrence of dipeptide EE indicates the significance of glutamic acid in intrinsically disordered regions. The role of glutamic acid in ordered and disordered proteins is discussed in a recent review by Uversky (2013).

Another interesting observation is the abundance of the dipeptide with glycine and proline in sequences of 100% predicted IDP regions. It should be noted that the proline is unable to occupy many of the main chain conformations easily whereas glycine can adopt due to its smaller size and hence proline can be considered as an opposite to glycine. The maximum number of occurrences of two residues with different properties may play a vital role in promoting disorder. The study by Campion et al., (2001) on ordered proteins shows that the dipeptide with glycine and proline has a minimum number of occurrences whereas it occurs more often in 100% predicted IDP regions.

**Figure 4:**
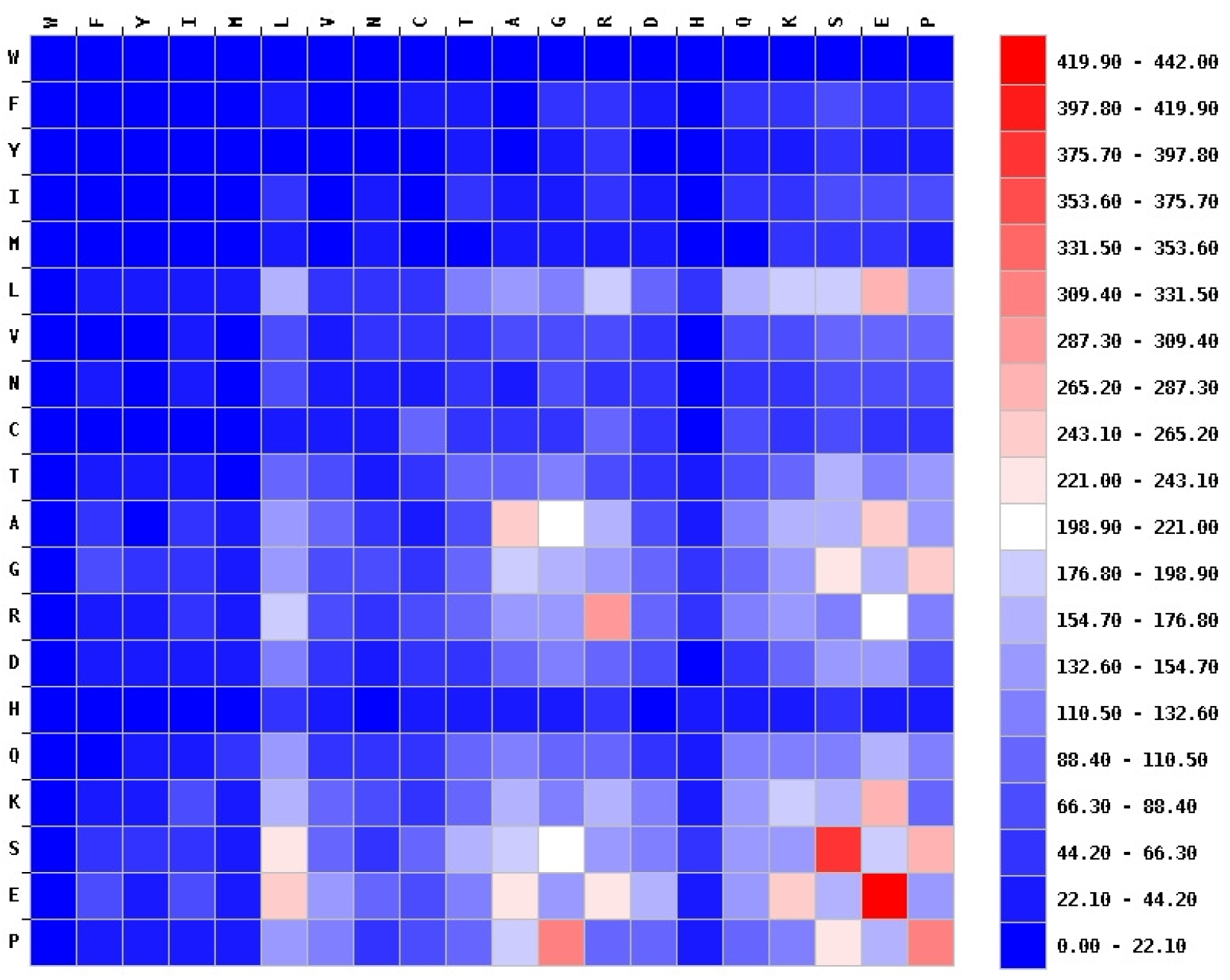
Pairwise residue-residue frequency distribution of 100% predicted IDP regions. The number of occurrences of dipeptides in 100% predicted IDPs are shown as grid diagram. The higher occurrences are represented by red color and lower occurrences as blue color. Whereas the dipeptide EE occurs maximum; SS, PP, PG, RR, AA, LE, AE and GP occur with high frequency. It can be seen that the first five order promoting (WFYIM) residues have a minimum number of occurrences as shown by blue grids.

### SEQUENCE FINGERPRINTING OF PREDICTED INTRINSIC DISORDER

Knowledge of how frequently two amino acids are separated along the sequence may help to understand the protein folding process and hence, we have computed the sequence separation of all possible pairs of amino acid residues in sequences of 100% predicted IDP regions and compared them with sequence separation values in proteins belonging to four major structural classes in SCOP database (Murzin et al., 1995). The correlation matrix between the number of occurrences of distance of separation between two amino acid residues along a sequence (sequence separation) of 100% predicted IDP regions with the occurrences of pair of amino acid residues at sequence separations of all alpha class of proteins is shown in Figure 5A. From the figure, it is observed that the distribution of cysteine residues along the sequence is completely different when compared to the 100% predicted IDP regions with sequences of various structural classes of proteins. Since cysteine is known to play an important role in protein folding and stability (Netto et al., 2007), our observation reveals that the absence of proper distribution of cysteine along the sequence may be a reason for intrinsic disorder. The similar trend is found for the correlation coefficient between the number of occurrences of a pair of amino acid residues at sequence separations of 100% predicted IDP regions and all beta, alpha+beta and alpha/beta classes of proteins which are shown in Figure 5 B, C and D.

**Figure 5A:**
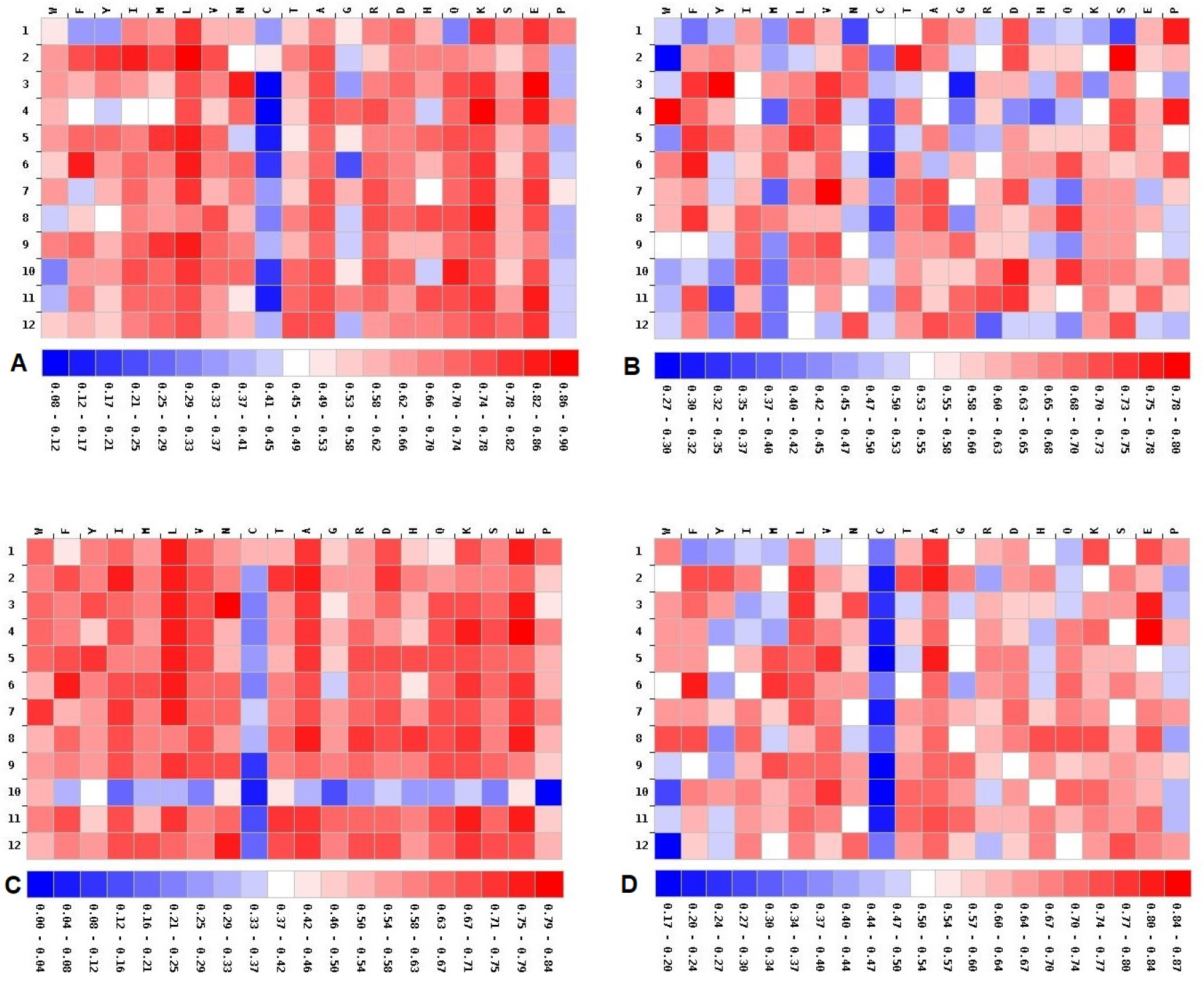
The correlation matrix with respect to 100% predicted IDPs and all alpha class proteins. The correlation matrix is shown between the numbers of occurrences of pair of amino acid residues upto 12 residue sequence separations (distance of separation between two amino acid residues along a sequence) of 100% predicted IDPs with the occurrences of pair of amino acid residues at similar sequence separations of all alpha class of proteins. The absence of proper distribution of cysteine along the sequence may be a reason for intrinsic disorder. Figure 5B: The correlation matrix with respect to 100% predicted IDPs and all beta class proteins. The correlation matrix is shown between the numbers of occurrences of pair of amino acid residues upto 12 residue sequence separations (distance of separation between two amino acid residues along a sequence) of 100% predicted IDPs with the occurrences of pair of amino acid residues at similar sequence separations of all beta class of proteins. The depletion in C is also seen in this comparison. Figure 5C: The correlation matrix with respect to 100% predicted IDPs and alpha+beta class proteins. The correlation matrix is shown between the numbers of occurrences of pair of amino acid residues upto 12 residue sequence separations (distance of separation between two amino acid residues along a sequence) of 100% predicted IDPs with the occurrences of pair of amino acid residues at similar sequence separations of alpha+beta class of proteins. Interestingly the residue pairs separated by distance ten are completely different in 100% predicted IDP regions and alpha+beta sequences. Figure 5D: The correlation matrix with respect to 100% predicted IDPs and alpha/beta class proteins. The correlation matrix is shown between the numbers of occurrences of pair of amino acid residues upto 12 residue sequence separations (distance of separation between two amino acid residues along a sequence) of 100% predicted IDPs with the occurrences of pair of amino acid residues at similar sequence separations of alpha/beta class of proteins. The distribution of proline along the sequence in 100% predicted IDPs differs much more than in the ordered proteins.

In all the depletion with pairs involving cysteine is seen. Interestingly, only in the comparison with alpha+beta class, the correlation for a sequence separation of ten residues is low (figure 5C). This means that the residue pairs separated by distance ten are completely different in 100% predicted IDP regions and alpha+beta sequences. From our analysis, it is also found especially for alpha/beta class comparison (Figure 5D) that the distribution of proline along the sequence in 100% predicted IDPs differs much more than in the ordered proteins.

### DISTINGUISHING PATTERNS FOUND BETWEEN DISORDERED AND GLOBULAR DATA SETS

As described in the Materials and Methods section, a total of 8784 possible pentapeptide patterns (4392 pentapeptides made up of 5 disorder promoting residues+4392 pentapeptides made up of 5 order promoting residues) were generated based on the previously described scale^38^. There are considerable number of occurrences of pentapeptide patterns made up of disorder promoting amino acid residues are found in both Disprot and globular proteins which is clearly reflected in the log-odds analysis (Table 2a,b). The difference between the sum of log-odds of the pentaXd sets and the pentaXo sets shows a remarkable pattern of being negative for the 100%IDP (−0.76 bits) and Disprot data (−1.49 bits) while being positive for the Globular protein data (3.79 bits). This means that it is the combination of occurrences of patterns of the pentaX sets comprising the disorder and order carries the distinguishing feature to differentiate the erroneous from the correct predictions rather than any one pattern by itself.

Interestingly, we have not found any matches while searching pentapeptide patterns comprising order promoting residues in the sequences of 100% predicted IDPs and Disprot database. This shows that there is no combination of top 3 order promoting residues occurring in the sequences of predicted and native intrinsically disordered regions. The pentapeptide patterns thus generated were also matched with sequences of the globular proteins dataset (Singh et al., 2012). The pentapeptide patterns comprising both order promoting and disorder promoting amino acid residues are found in globular proteins. Further, we made a pattern search with pentapeptide patterns comprising disorder promoting residues in the sequences of alpha and beta transmembrane regions (residues with a regular DSSP (Kabsch & Sander, 1983) assigned secondary structure like H and E) compiled by us for our previous work (Saravanan & Krishnaswamy, 2015) and found no matches.

**Table 2a.**
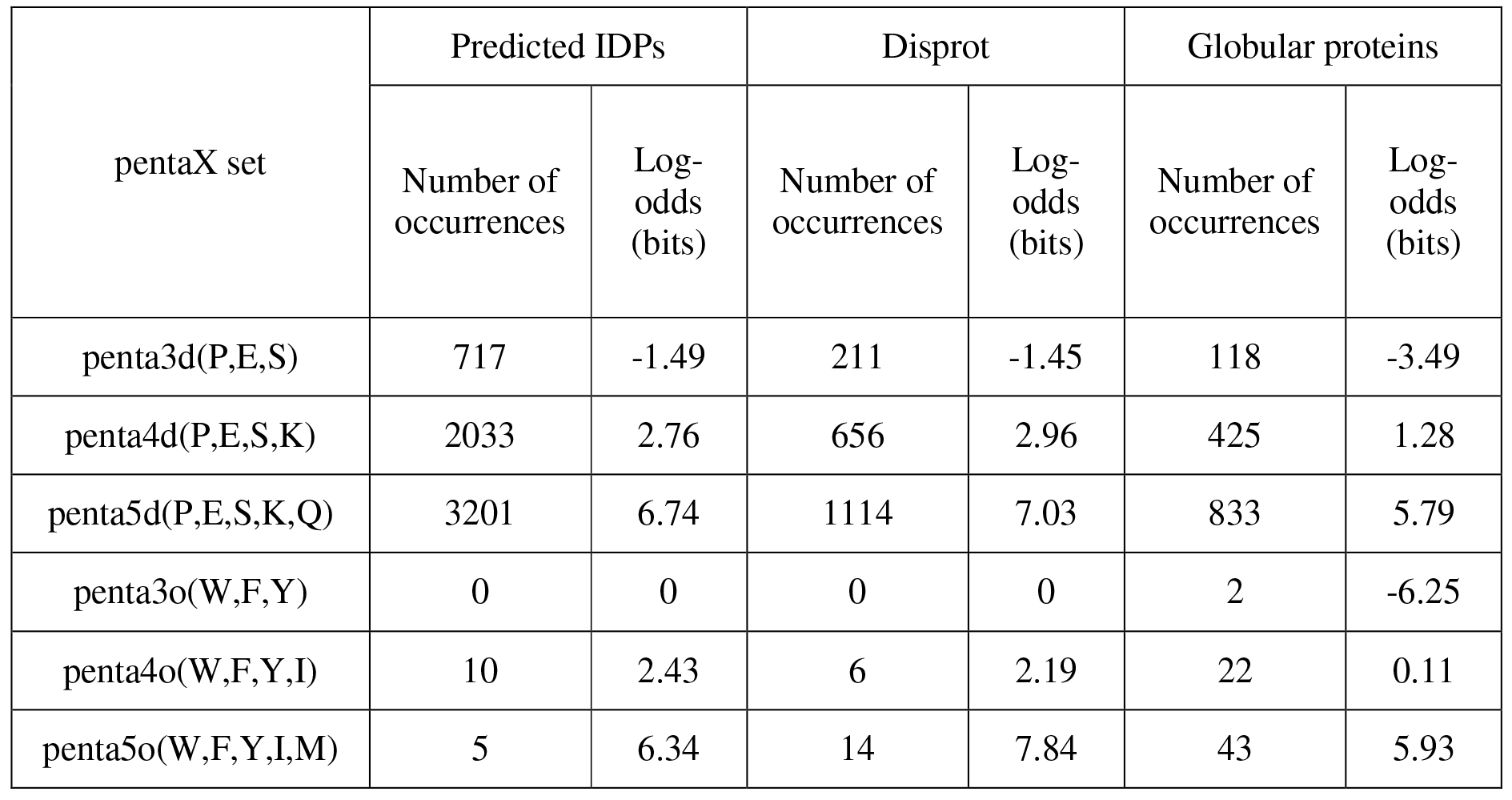
Occurence of pentapeptide combinations in predicted IDPs, native intrinsic disordered and globular protein sequences.

**Table 2b.** Log odds (bits) difference in occurrence between disordered and ordered sets.

| pentaX set | 100% IDP | Disprot | Globular |
| --- | --- | --- | --- |
| penta3d(P,E,S) | -1.49 | -1.45 | -3.49 |
| penta4d(P,E,S,K) | 2.76 | 2.96 | 1.28 |
| penta5d(P,E,S,K,Q) | 6.74 | 7.03 | 5.79 |
| Sum of pentaXd sets | 8.01 | 8.54 | 3.58 |
| penta3o(W,F,Y) | 0 | 0 | -6.25 |
| penta4o(W,F,Y,I) | 2.43 | 2.19 | 0.11 |
| penta5o(W,F,Y,I,M) | 6.34 | 7.84 | 5.93 |
| Sum pentaXo sets | 8.77 | 10.03 | -0.21 |
| **Sum (pentaXd) –Sum (pentaXo)** | **-0.76** | **-1.49** | **3.79** |

Since the above generated pentapeptides contains at least three disorder promoting residues, the patterns with repeated amino acid residues like PPP or EEE were not used in the above pattern search analysis and hence we have made such a pattern search and presented the results in Table 3. The log-odds of occurrences of patterns clearly reveal that the abundance of pattern of proline residues (PPP and PPPP) is higher in intrinsic disordered regions than the globular proteins with regular secondary structure. Consistently the log-odds of occurrences of the repeating pattern of residues (P or E or S) is more negative for the Globular protein dataset than the 100%IDP dataset or Disprot dataset (eg −1.15 bits for PPP in 100%IDP, −6.89 bits in Disprot whereas −9.32 bits in Globular). This arises from the occurrence of these distinguishing patterns of repeats in the Globular proteins being much less than in the random case. The difference in the sum of the log odds between the 100%IDP and Globular sets is 62.59 bits while the difference between the Disprot and Globular sets is 21.61 bits. The large positive values suggest significant information content in the difference of the occurrence of repeated patterns of P or E or S below the pentapeptide level between the Globular proteins and the Intrinsically Disordered Proteins. The tri and tetra repeats of P or E or S provide a distinguishing feature to detect the erroneous and correct predictions.

**Table 3.** Occurrences of patterns comprising repeated disorder promoting amino acid residues.

| Pattern | Number of occurrences |  |  | Log-odds of occurrences (bits) |  |  |  |  |
| --- | --- | --- | --- | --- | --- | --- | --- | --- |
|  | Predicted IDP | DisProt | Globular Proteins | Predicted IDP | DisProt | Globular Proteins | Diff Predicted IDP-Glob | Diff DisProt-Glob |
| PPP | 758 | 235 | 164 | -1.15 | -6.89 | -9.32 | 8.17 | 2.43 |
| PPPP | 211 | 56 | 18 | -0.34 | -9.25 | -12.78 | 12.44 | 3.53 |
| EEE | 947 | 451 | 472 | -1.21 | -6.38 | -10.03 | 8.82 | 3.65 |
| EEEE | 237 | 122 | 32 | -0.7 | -8.5 | -13.49 | 12.79 | 4.99 |
| SSS | 805 | 324 | 393 | -1.47 | -6.74 | -9.79 | 8.32 | 3.05 |
| SSSS | 163 | 66 | 24 | -1.26 | -9.35 | -13.31 | 12.05 | 3.96 |
| Sum log-odds |  |  |  | -6.13 | -47.11 | -68.72 | **62.59** | **21.61** |

While searching for palindromes in sequences of 100% PID regions, we found 1819 palindromes with 3 residues, 184 palindromes with 4 residues, 227 palindromes with 5 residues, 26 palindromes with 6 residues and 27 palindromes with 7 residues. The presence of such number of palindromes may be due to the low complexity nature of the 100% predicted IDP sequences. The percentage of occurrences of short hydrophobic stretches in predicted and native IDPs reveal the absence of significant differences as shown in Figure 6. It is possible to incorporate the information regarding the slightly higher occurrence of 3 residue long hydrophobic stretches in the native IDPs, and the contrasting marginally more occurrence albeit not substantial of the 4,5,6 and 7 residue stretches in the predicted IDPs for use in a program to identify erroneous from correct predictions.

**Figure 6.**
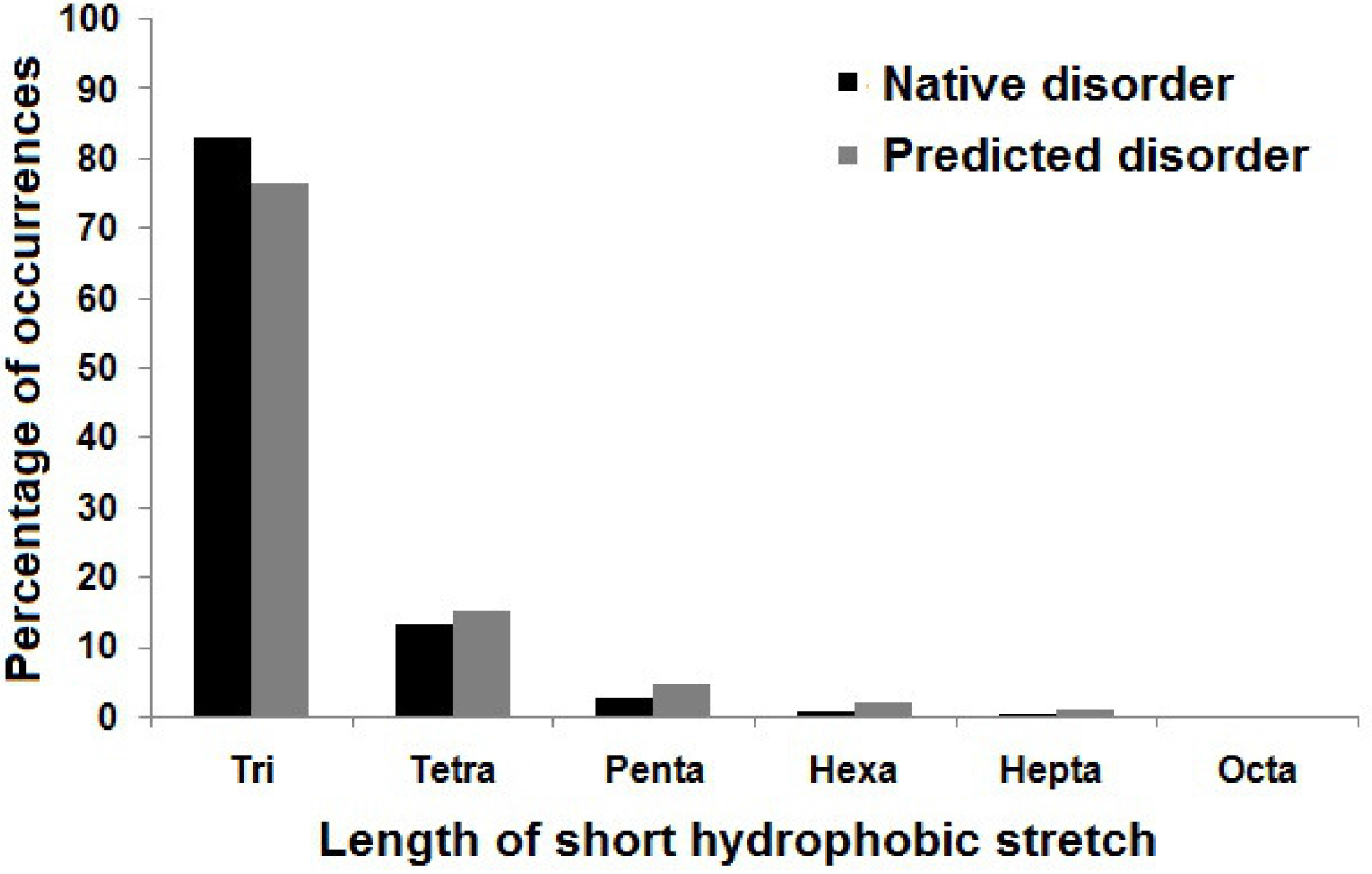
Percentage of occurrences of short hydrophobic stretches in sequences of predicted and native IDPs. The length of the hydrophobic stretches in predicted and native IDPs are grouped as trimers (residue length of 3), tetramers (residue length of four), pentamers (residue length of 5), hexamers (residue length of 6), heptamers (residue length of 7) and octamers (residue length of 8). It may be significant that the occurrence of 3 residue long hydrophobic stretches are more in the native IDPs, the 4,5,6 and 7 residue stretches occur slightly more in the predicted IDPs.

### SECONDARY STRUCTURE PREDICTION IN 100% PREDICTED IDP SEQUENCES

The percentage of predicted secondary structural elements such as helix, extended and coil respectively occurring in 100% predicted IDPs is shown in figure 7. The results of consensus secondary structure prediction are given as a single text file in **Supplementary Material 1**. From these results, it is noted that the 64% of residues in the sequences of 100% predicted IDP regions are predicted as coils and 32.4% of residues are predicted as helices. Only 3.6% of residues are predicted as extended (e.g. β-sheet) regions. It is quite likely that a significant fraction of the predicted helix regions actually forms helix upon binding to partners. 32.4% of residues are predicted as helices and this may be due to the classification of amino acid residues based on their secondary structure forming propensity in the secondary structure prediction algorithms. From our analysis, we suggest that the secondary structure prediction methods can also be used as distinguishing factor in disorder predictions by making suitable changes in the secondary structure prediction algorithm and by classifying residues as order and disorder promoters.

**Figure 7:**
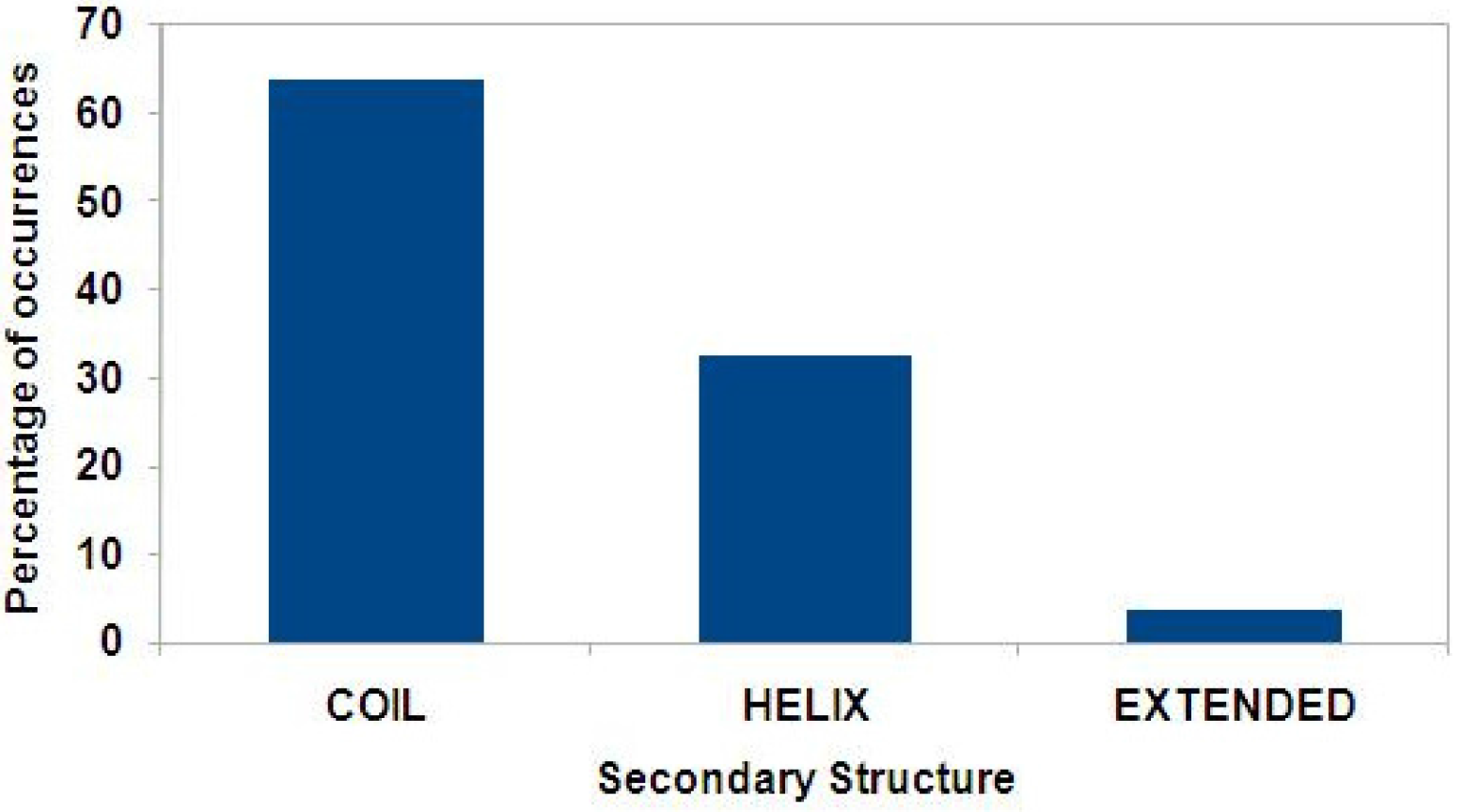
Secondary structure prediction of 100% predicted IDPs. The percentage of occurrences of amino acid residues in secondary structural elements like helix, extended and coil respectively in 100% predicted IDRs. It is seen 64% of residues in the sequences of 100% predicted IDP regions are predicted as coils and 32.4% of residues are predicted as helices. Only 3.6% of residues are predicted as extended (e.g. β-sheet) regions.

## DISCUSSION

Predicting tertiary structure of proteins is still a relatively unsolved problem in computational molecular biology. In this proteomics and structural genomics era, identifying proteins with intrinsic disorder is another challenging problem (Linding et al., 2003). Similarity and dissimilarity analysis have been reported to understand the characteristics of ordered and natively disordered regions(Romero et al., 2001). Weakness in prediction of IDPs and IDRs are due to various factors like amino acid composition, sequence length, sequence repeats and sequence entropy etc., (Kurup, Dunker & Krishnaswamy, 2013). Several disorder predictors have been developed by using different theories such as pseudo amino acid composition, neural networks and machine learning respectively which are discussed in a review by Dosztanyi et al.,(2007). Since, the accuracy of disordered predictors has reached a reasonable accuracy; it was of interest to analyze the predicted IDP regions by using computational approaches like amino acid composition, dipeptide analysis, sequence separation of pair of amino acids, pattern searching and secondary structure prediction.

Our sequence fingerprinting of 100% predicted IDP regions is in agreement with the amino acid scale developed by Campen et al., (2008) based on the native disordered proteins. Differences between the composition of residues like C and R is observed in native intrinsic disordered regions and 100% PID regions. Dipeptide analysis of 100% predicted IDP regions reveal the abundance of two amino acids G and P with opposite properties in adopting an ordered structure. The frequency of occurrences of dipeptide GP is high in 100% predicted IDP regions which may promote intrinsic order whereas it is low in the case of ordered regions (Campion et al., 2001). The distribution and placement of cysteine residues along the polypeptide chain is different while comparing sequence separation of 100% predicted IDP regions and sequences of proteins in four major structural classes.

Pattern searching on 100% predicted IDP regions reveal several interesting observation like the lack of occurrence of pentapeptide patterns with top three order promoting amino acid residues in the sequences of 100% predicted and native intrinsically disordered proteins. The results of pattern search analysis reveal that no combination of top 3 order promoting amino acid residues is preferred in the sequences of predicted and native IDP regions. The pattern with repeated proline residues is abundant in intrinsic disordered regions than the globular proteins with regular secondary structure. No hits are obtained while searching for pentapeptide patterns with disorder promoting residues in the sequences of transmembrane regions. From our secondary structure prediction results, it is noted that 64% of residues in 100% predicted IDP regions is predicted as coils. Only a small percentage of residues are predicted as extended regions. The distinguishing features presented based on both residue basis (amino acid based) and segment basis (sequence fragments) in this paper may help to understand the role of sequence fingerprints in detecting erroneous from correct predictions of IDPs and IDP regions. We also suggest that the differences observed between native disorder and predicted disorder can be used to improve the accuracy of disorder predictors.

## ACKNOWLEDGMENTS

KMS is supported by a National Post-Doctoral Fellowship (File No: PDF/2015/000276) by Science Engineering Research Board (SERB), Government of India, under the mentorship of Dr. P. Karthe, Professor of Crystallography & Biophysics at the University of Madras.

## AUTHOR CONTRIBUTIONS

Experiments were designed by SK and AKD, carried out by KMS, data was analyzed by KMS, AKD and SK. The paper was written by KMS, AKD and SK.

## SUPPORTING INFORMATION

Supplementary Material 1: Composition differences in sequences of predicted and native intrinsic disorder.

Supplementary Material 2: Consensus secondary structure prediction of intrinsic disordered regions.

## REFERENCES

Banerjee, S., Chakraborty, S., & De, R. K. (2017). Deciphering the cause of evolutionary variance within intrinsically disordered regions in human proteins. J. Biomol. Struct. Dyn. 35, 233–249.

Bellay, J., Han, S., Michaut, M., Kim, T., Costanzo, M., Andrews, B. J., Boone, C., Bader, G. D., Myers, C. L., & Kim, P. M. (2011). Bringing order to protein disorder through comparative genomics and genetic interactions. Genome Biol. 12, R14.

Brown, C. J., Johnson, A. K., Dunker, A. K., & Daughdrill, G. W. (2011). Evolution and disorder. Curr. Opin. Struct. Biol. 21, 441–446.

Brown, C. J., Takayama, S., Campen, A. M., Vise, P., Marshall, T. W., Oldfield, C. J., Williams, C. J., & Dunker, A. K. (2002). Evolutionary rate heterogeneity in proteins with long disordered regions. J. Mol. Evol. 55, 104–110.

Campen, A., Williams, R. M., Brown, C. J., Meng, J., Uversky, V. N., & Dunker, A. K. (2008). TOP-IDP-scale: a new amino acid scale measuring propensity for intrinsic disorder. Protein Pept. Lett. 15, 956–963.

Campion, S. R., Ameen, A. S., Lai, L., King, J. M., & Munzenmaier, T. N. (2001). Dipeptide frequency/bias analysis identifies conserved sites of non-randomness shared by cysteine-rich motifs. Proteins 44, 321–328.

Chen, J. W., Romero, P., Uversky, V. N., & Dunker, A. K. (2006). Conservation of intrinsic disorder in protein domains and families: I. A database of conserved predicted disordered regions. J. Proteome Res. 5, 879–887.

Coelho Ribeiro Mde, L., Espinosa, J., Islam, S., Martinez, O., Thanki, J. J., Mazariegos, S., Nguyen, T., Larina, M., Xue, B., & Uversky, V. N. (2013). Malleable ribonucleoprotein machine: protein intrinsic disorder in the Saccharomyces cerevisiae spliceosome. Peer J. 12, e2.

Di Domenico, T., Walsh, I., Martin, A. J., & Tosatto, S. C. (2012). MobiDB: a comprehensive database of intrinsic protein disorder annotations. Bioinformatics 28, 2080–2081.

Dosztányi, Z., Chen, J., Dunker, A. K., Simon, I., & Tompa, P. (2006). Disorder and sequence repeats in hub proteins and their implications for network evolution. J. Proteome Res. 5, 2985–2995.

Dosztányi, Z., Csizmok, V., Tompa, P., & Simon, I. (2005). IUPred: web server for the prediction of intrinsically unstructured regions of proteins based on estimated energy content. Bioinformatics 21, 3433–3434.

Dosztányi, Z., Sándor, M., Tompa, P., & Simon, I. (2007). Prediction of protein disorder at the domain level. Curr. Prot. Pept. Sci. 8, 161–171.

Dunker, A. K., & Oldfield, C. J. (2015). Back to the future: Nuclear magnetic resonance and bioinformatics studies on intrinsically disordered proteins. Adv. Exp. Med. Biol. 870, 1–34.

Dunker, A. K., Lawson, J. D., Brown, C. J., Williams, R. M., Romero, P., Oh, J. S., & Obradovic, Z. (2001). Intrinsically disordered protein. J. Mol. Graph. Model. 19, 26–59.

Dunker, A. K., Obradovic, Z., Romero, P., Garner, E. C., & Brown, C. J. (2000). Intrinsic protein disorder in complete genomes. Genome Inform. 161–171.

Dyson, H. J., & Wright, P. E. (2005). Intrinsically unstructured proteins and their functions. Nature Rev. Mol. Cell Biol. 6, 197–208.

Forman-Kay, J. D., & Mittag, T. (2013). From sequence and forces to structure, function, and evolution of intrinsically disordered proteins. Structure 21, 1492–1499.

Fraga, H., Grana-Montes, R., Illa, R., Coveleda, G., & Ventura, S. (2014). Association between foldability and aggregation propensity in small disulfide-rich proteins. Antioxid. Redox Signal. 21, 368–383.

Frishman, D., & Argos, P. (1997). Seventy-five percent accuracy in protein secondary structure prediction. Proteins 27, 329–335.

Fukuchi, S., Homma, K., Minezaki, Y., & Nishikawa, K. (2006). Intrinsically disordered loops inserted into the structural domains of human proteins. J. Mol. Biol. 355, 845–857.

Fukuchi, T., Amemiya, T., Sakamoto, S., Nobe, Y., Hosoda, K., Kado, Y., Murakami, S. D., Koike, R., Hiroaki, H., & Ota, M. (2014). IDEAL in 2014 illustrates interaction networks composed of intrinsically disordered proteins and their binding partners. Nucleic Acids Res. 42, D320–D325.

Garnier, J., Gibrat, J. F., & Robson, B. (1996). GOR method for predicting protein secondary structure from amino acid sequence. Methods Enzymol. 266, 540–553.

Guermeur, Y., Geourjon, C., Gallinari, P., & Del, G. (1999). Improved performance in protein secondary structure prediction by inhomogeneous score combination. Bioinformatics 15, 413–421.

Gunasekaran, K., Tsai, C. J., Kumar, S., Zanuy, D., & Nussinov, R. (2003). Extended disordered proteins: targeting function with less scaffold. Trends Biochem. Sci. 28, 81–85.

Gupta, A., Deshpande, A., Amburi, J. K., Sabarinathan, R., Senthilkumar, R., & Sekar, K. (2009). CSSP (consensus secondary structure prediction): a web-based server for structural biologists. J. App. Cryst. 42, 336–338.

Harihar, B., & Selvaraj, S. (2009). Refinement of the long-range order parameter in predicting folding rates of two-state proteins. Biopolymers 91, 928–935.

He, B., Wang, K., Liu, Y., Xue, B., Uversky, V. N., & Dunker, A. K. (2009). Predicting intrinsic disorder in proteins: an overview. Cell Res. 19, 929–949.

Ishida, T., & Kinoshita, K. (2007). PrDOS: prediction of disordered protein regions from amino acid sequence. Nucleic Acids Res. 35, 460–464.

Kabsch, W., & Sander, C. (1983). Dictionary of protein secondary structure: pattern recognition of hydrogen-bonded and geometrical features. Biopolymers 22, 2577–2637.

King, R. D., & Sternberg, M. J. (1996). Identification and application of the concepts important for accurate and reliable protein secondary structure prediction. Protein Sci. 5, 2298–2310.

Korneta, I., Bujnicki, J. M. (2012). Intrinsic disorder in the human spliceosomal proteome. PLoS Comput. Biol. 8, e1002641.

Kosol, S., Contreras-Martos, S., Cedeño, C., & Tompa, P. (2013). Structural characterization of intrinsically disordered proteins by NMR spectroscopy. Molecules 18, 10802–10828.

Kurup, K., Dunker, A. K., & Krishnaswamy, S. (2013). Functional fragments of disorder in outer membrane beta barrel proteins. Intrinsically Disordered Proteins 1, 45–55.

Lee, K. H., Zhang, P., Kim, H. J., Mitrea, D. M., Sarkar, M., Freibaum, B. D., Cika, J., Coughlin, M., Messing, J., Molliex, A., Maxwell, B. A., Kim, N. C., Temirov, J., Moore, J., Kolaitis, R. M., Shaw, T. I., Bai, B., Peng, J., Kriwacki, R. W., & Taylor, J. P. (2016). C9orf72 dipeptide repeats impair the assembly, dynamics and function of membrane less organelles. Cell 167, 774–788.

Linding, R., Jensen, L. J., Diella, F., Bork, P., Gibson, T. J., & Russell, R. B. (2003). Protein disorder prediction: implications for structural proteomics. Structure 11, 1453–1459.

Lobley, A., Swindells, M. B., Orengo, C. A., & Jones, D. T. (2007). Inferring function using patterns of native disorder in proteins. PLoS Comput. Biol. 3, e162.

McGuffin, L. J., Bryson, K., & Jones, D. T. (2000). The PSIPRED protein structure prediction server. Bioinformatics 16, 404–405.

Meszaros, B., Dosztanyi, Z., & Simon, I. (2012). Disordered binding regions and linear motifs - bridging the gap between two models of molecular recognition. Plos One 7, e46829.

Minde, D. P., Dunker, A. K., & Lilley, K. S. (2017). Time, space and disorder in the expanding proteome universe. Proteomics 17, 7.

Monastyrskyy, B., Kryshtafovych, A., Moult, J., Tramontano, A., & Fidelis, K. (2014). Assessment of protein disorder region predictions in CASP10. Proteins 82, 127–137.

Murzin, A. G., Brenner, S. E., Hubbard, T., & Chothia, C. (1995). SCOP: a structural classification of proteins database for the investigation of sequences and structures. J. Mol. Biol. 247, 536–540.

Netto LES, de Oliveira, M. A., Monteiro, G., Demasi, A. P. D., Cussiol, J. R. R., Discola, K. F., & Horta, B. B. (2007). Reactive cysteine in proteins: protein folding, antioxidant defense, redox signalling and more. Comp. Biochem. Physiol. 146, 180–193.

Oates, M. E., Romero, P., Ishida, T., Ghalwash, M., Mizianty, M. J., Xue, B., & Gough, J. (2013). D2P2: database of disordered protein predictions. Nucleic Acids Res. 41, D508–D516.

Obradovic, Z., Peng, K., Vucetic, S., Radivojac, P., & Dunker, A. K. (2005). Exploiting heterogeneous sequence properties improves prediction of protein disorder. Proteins 61, 176–182.

Obradovic, Z., Peng, K., Vucetic, S., Radivojac, P., Brown, C. J., & Dunker, A. K. (2003). Predicting intrinsic disorder from amino acid sequence. Proteins 53, 566–572.

Oldfield, C. J., & Dunker, A. K. (2014). Intrinsically Disordered Proteins and Intrinsically Disordered Protein Regions. Ann. Rev. Biochem. 83, 553–584.

Peng, Z. L., & Krugan, L. (2012). Comprehensive comparative assessment of in silico predictors of disordered regions. Curr. Protein Pept. Sci. 13, 6–18.

Peng, Z., Oldfield, C. J., Xue, B., Mizinty, M. J., Dunker, A. K., Kurgan, L., & Uversky, V. N. (2014). A creature with a hundred waggly tails: intrinsically disordered proteins in the ribosome. Cell Mol. Life Sci. 71, 1477–1504.

Piovesan, D., Tabaro, F., Micetic, I., Necci, M., Quaglia, F., Oldfield, C. J., Aspromonte, C., Davey, N. E., Davidovic, R., Dosztanyi, Z., Elofsson, A., Gasparini, A., Hatos, A., Kajava, A. V., Kalmar, L., Leonardi, E., Lazar, T., Macedo-Ribeiro, S., Macossay-Castillo, M., Meszaros, A., Minervini, G., Murvai, N., Pujols, J., Roche, D. B., Salladini, E., Schad, E., Schramm, A., Szabo, B., Tantos, A., Tonello, F., Tsirigos, K. D., Veljkovic, N., Ventura, S., Vranken, W., Warholm, P., Uversky, V. N., Dunker, A. K., Longhi, S., Tompa, P., & Tosatto, S. C. E. (2017). Disprot 7.0: a major update of the database of disordered proteins. Nucleic Acids Res 45, D219–D227.

Potenza, E., Domenico, T. D., Walsh, I., & Tosatto, S. C. (2015). MobiDB 2.0: an improved database of intrinsically disordered and mobile proteins. Nucleic Acids Res. 43, D315–320.

Radivojac, P., Iakoucheva, L. M., Oldfield, C. J., Obradovic, Z., Uversky, V. N., & Dunker, A. K. (2007). Intrinsic disorder and functional proteomics. Biophys. J. 92, 1439–1456.

Romero, P., Obradovic, Z., Kisinger, K., Villafranca, J. E., & Dunker, A. K. (1997). Identifying disordered regions in proteins from amino acid sequence, Int. Conf. Neural Net. 1, 90–95.

Romero, P., Obradovic, Z., Li, X., Garner, E. C., Brown, C. J., & Dunker, A. K. (2001). Sequence complexity of disordered protein. Proteins 42, 38–48.

Rost, B., & Eyrich, V. A. (2001). EVA: large scale analysis of secondary structure prediction. Proteins 45, 192–199.

Saravanan, K. M., & Krishnaswamy, S. (2015). Analysis of dihedral angle preferences for alanine and glycine residues in alpha and beta transmembrane regions. J. Biomol. Struct. Dyn. 33, 5525–562.

Saravanan, K. M., & Selvaraj, S. (2017). Dihedral angle preferences of amino acid residues forming non-local interactions in proteins. J. Biol. Phys. 43, 265–278.

Schlessinger, A., Schaefer, C., Vicedo, E., Schmidberger, M., Punta, M., & Rost, B. (2011). Protein disorder – a breakthrough invention of evolution? Curr. Opin. Struct. Biol. 21, 412–418.

Schmidt, H. B., & Görlich, D. (2016). Transport Selectivity of Nuclear Pores, Phase Separation, and Membraneless Organelles. Trends Biochem. Sci. 41, 46–61.

Shimizu, K., Hirose, S., & Noguchi, T. (2007). POODLE-S: web application for predicting protein disorder by using physicochemical features and reduced amino acid set of a position-specific scoring matrix. Bioinformatics 23, 2337–2338.

Sickmeier, M., Hamilton, J. A., LeGall, T., Vacic, V., Cortese, M. S., Tantos, A., Szabo, B., Tompa, P., Chen, J., Uversky, V. N., Obradovic, Z., & Dunker, A. K. (2007). DisProt: the Database of Disordered Proteins. Nucleic Acids Res. 35, D786–793.

Simon, M., & Hancock, J. M. (2009). Tandem and cryptic amino acid repeats accumulate in disordered regions of proteins. Genome Biol. 10, R59.

Singh, H., Chauhan, J. S., Gromiha, M. M., & Raghava, G. P. (2012). ccPDB: compilation and creation of data sets from Protein Data Bank. Nucleic Acids Res. 40, 486–489.

Szalkowski, A. M., & Anisimova, M. (2011). Markov models of amino acid substitution to study proteins with intrinsically disordered regions. PLoS One 6, e20488.

Tompa, P. (2002). Intrinsically unstructured proteins. Trends Biochem. Sci. 2002, 27, 527–533.

Tompa, P., Fuxreiter, M., Oldfield, C. J., Simon, I., Dunker, A. K., & Uversky, V. N. (2009). Close encounters of the third kind: disordered domains and the interactions of proteins. Bioessays 31, 328–335.

Toretsky, J. A., & Wright, P. E. (2014). Assemblages: functional units formed by cellular phase separation. J. Cell. Biol. 206, 579–588.

Tóth-Petróczy, A., Oldfield, C. J., Simon, I., Takagi, Y., Dunker, A. K., Uversky, V. N., & Fuxreiter, M. (2008). Malleable machines in transcription regulation: the mediator complex. PLoS Comput. Biol. 4, e1000243.

Uversky, V. N. (2002). Natively unfolded proteins: a point where biology waits for physics. Protein Sci. 11, 739–756.

Uversky, V. N. (2013). The alphabet of intrinsic disorder: II. Various roles of glutamic acid in ordered and intrinsically disordered proteins. Intrinsically Disordered Proteins 1, 18–40.

Uversky, V. N., Kuznetsova, I. M., Turoverov, K. K., & Zaslavsky, B. (2015). Intrinsically disordered proteins as crucial constituents of cellular aqueous two phase systems and coacervates. FEBS Lett. 589, 15–22.

Vacic, V., Uversky, V. N., Dunker, A. K., & Lonardi, S. (2007). Composition Profiler: A tool for discovery and visualization of amino acid composition differences. BMC Bioinformatics 8, 211.

Varadi, M., Kosol, S., Lebrun, P., Valentini, E., Blackledge, M., Dunker, A. K., & Tompa, P. (2014). pE-DB: a database of structural ensembles of intrinsically disordered and of unfolded proteins. Nucleic Acids Res. 42, D326–D335.

Walsh, I., Martin, A.J., Di Domenico, T., & Tosatto, S. C. (2012). ESpritz: accurate and fast prediction of protein disorder. Bioinformatics 28, 503–509.

Wang, G., & Dunbrack, R. L. (2003). PISCES: a protein sequence culling server. Bioinformatics 19, 1589–1591.

Ward, J. J., Sodhi, J. S., McGuffin, L. J., Buxton, B. F., & Jones, D. T. (2004). Prediction and functional analysis of native disorder in proteins from the three kingdoms of life. J. Mol. Biol. 337, 635–645.

Weathers, E. A., Paulaitis, M. E., Woolf, T. B., & Hoh, J. H. (2004). Reduced amino acid alphabet is sufficient to accurately recognize intrinsically disordered protein. FEBS Lett. 576, 348–352.

Williams, R. W., Xue, B., Uversky, V. N., & Dunker, A. K. (2013). Distribution and cluster analysis of predicted intrinsically disordered protein Pfam domains. Intrinsically disordered proteins 1, e24848.

Wright, P. E., & Dyson, H. J. (1999). Intrinsically unstructured proteins: re-assessing the protein structure-function paradigm. J. Mol. Biol. 293, 321–331.

Wu, H., & Fuxreiter, M. (2016). The structure and dynamics of higher-order assemblies: amyloids, signalosomes, and granules. Cell 165, 1055–1066.

Xue, B., Dunbrack, R. L., Williams, R. W., Dunker, A. K., & Uversky, V. N. (2010). PONDR-FIT: a meta-predictor of intrinsically disordered amino acids. Biochim. Biophys. Acta. 1804, 996–1010.

Xue, B., Dunker, A. K., & Uversky, V. N. (2012). Orderly order in protein intrinsic disorder distribution: disorder in 3500 proteomes from viruses and the three domains of life. J. Biomol. Struct. Dyn. 30, 137–149.

Xue, B., Oldfield, C. J., Dunker, A. K., & Uversky, V. N. (2009). CDF it all: Consensus prediction of intrinsically disordered proteins based on various cumulative distribution functions. FEBS Lett. 583, 1469–1474.

Xue, B., Romero, P. R., Noutsou, M., Maurice, M. M., Rüdiger, S. G., William, A. M. Jr., Mizianty, M. J., Kurgan, L., Uversky, V. N., & Dunker, A. K. (2013). Stochastic machines as a colocalization mechanism for scaffold protein function. FEBS Lett. 587, 1587–1591.

Yan, J., Dunker, A. K., Uversky, V. N., & Kurgan, L. (2016). Molecular recognition features (MoRFs) in three domains of life. Mol. Biosyst. 12, 697–710.

Yeon, J. H., Heinkel, F., Sung, M., Na, D., Gsponer, J. (2016). Systems wide identification of cis-regulatory elements in proteins. Cell Syst. 2, 89–100.

